# Phenotypic cross-species conservation and cross-generation directionality switching in epigenetic inheritance

**DOI:** 10.1101/2022.09.30.510079

**Authors:** Ameek Bhalla, Abhay Sharma

## Abstract

Evidence supporting non-DNA sequence-based inheritance in animals has increasingly been described in recent years, often under short-term, intergenerational inheritance or longer, transgenerational epigenetic inheritance (TEI). Existence of the latter, a stronger indicator of germline transmission, though established in invertebrates remains controversial in mammals due to inherent confounding factors. Besides evolutionary conservation, physiological implications of TEI also remain unclear. Leveraging invertebrate evidence of TEI to assess possible instances in mammals, and dissecting already described models to gain further insights are suggested approaches to address uncertainties in non-genetic inheritance. Here, in an unbiased approach, we compared existing transcriptomic data associated with so far available *Drosophila* models of inter- and trans-, and rodent models of inter-generational inheritance, observed phenotypic cross-species conservation and cross-generation directionality shift therein, and confirmed these observations experimentally in flies. Specifically, previous models of cold and diet induced inheritance in both flies and mice were commonly associated with altered regulation of proteolysis genes. Besides, fly TEI models were in general characterized by opposite phenotypic regulation between inter- and trans-generational offspring. As insulin producing cell (IPC) ablation was also associated with proteolysis gene dysregulation in one of the mouse models, we opted to use genetic ablation of IPCs in flies for the experimental confirmation. Remarkably, the ablation led to transcriptomic alterations across multiple generations, with dysregulated genes showing proteolysis enrichment. Similarly, phenotypic directionality changed in the opposite direction in transgenerational offspring of IPC ablated ancestors. These results support mammalian existence, and physiologically adaptive and maladaptive consequences of germline mediated epigenetic inheritance.

## Introduction

Non-DNA sequence-based inheritance of phenotypic traits across organismal generations, with factors like DNA methylation, histone modifications, and RNAs potentially carrying the heritable information, is a subject of immense current interest (*1–3*). True epigenetic inheritance would contradict the neo-Darwinian principle restricting the source of heredity to DNA sequence alone in general, suggest violation of Weismann barrier prohibiting soma-to-germline communication in animals where the two lineages separate early in development, and require mechanisms allowing germline transmission of epigenetic signals through gametogenesis, fertilization, and embryonic development, against the odds of chromatin reorganization and extensive epigenetic reprogramming in animal species (*1–3*). Affecting these fundamental concepts in biology, epigenetic inheritance could have intriguing implications for ecology, evolution, and human health and disease. Notably, evidence supporting non-genetic inheritance in animals has increasingly been described in recent years, mostly under short-term, intergenerational inheritance or longer, transgenerational epigenetic inheritance (TEI). Though both exhibit common features, the former is deficient in excluding the possibility that appearance of a phenotype – in F1 generation if the founder is a male, and up to F2 generation if the founder is a gestating female, in mammals, for example – might have resulted from a direct effect of the trigger rather than a heritable epigenetic factor (*1–3*). Evidence of TEI, which signifies survival of epigenetic information beyond intergenerational duration and hence suggests involvement of germline transmission, though well documented in invertebrate models, remains scanty and controversial in mammals. The bottleneck chiefly stems from difficulties in eliminating confounding influence of various factors such as maternal contribution, *in utero* and postnatal effects, and seminal fluid components, as also DNA sequence-based effects, in rat models that lack inbred stocks, for example (*1, 2*). Paternal exposure models with patrilineal inheritance are considered less fraught with confounding elements and hence a focus of current investigations (*2, 3*). Besides evolutionary conservation, physiological underpinnings in TEI also remain unclear. Whether non-genetic inheritance entails unintentional consequences or adaptive responses of ancestral environmental exposure is yet to be understood (*2, 3*).

It has been recently suggested that stronger evidence of TEI in invertebrate models may be leveraged to assess possible instances of the same in mammals, despite likely cross-species differences in the underlying mechanisms (*2*). Further dissecting the existing models of non-genetic inheritance in general is another recently suggested approach for gaining functional insights (*3*). The usefulness of these approaches is indeed supported by preliminary analysis comparing transcriptomic and metabolic responses in non-genetic inheritance (*4, 5*). As organisms may differ in their armamentarium of epigenetic mechanisms – with DNA methylation, for example, not considered a major, global epigenetic factor in *Drosophila melanogaster*, unlike vertebrates – regulation of gene expression, which various epigenetic modifications functionally converge on, may provide a means for investigating cross-species evolutionary conservation in non-genetic inheritance. Similarly, transcriptomic alterations may represent phenotypic changes at a comprehensive molecular level and hence prove informative in understanding epigenetic inheritance in terms of unintentional, or adaptive or maladaptive effects. Given these considerations, we first examine here existing transcriptomic data related to so far available paternal exposure and patrilineal *Drosophila* models of inter- and trans-generational, and mouse models of inter-generational inheritance. This data analysis suggests that transcriptomic responses are interchangeable between comparable fly and mouse models, and phenotypic directionality is changed along inter- to trans-generational progression in *Drosophila.* We next confirm these observations experimentally in flies.

## Results and Discussion

We first assembled a compendium of transcriptome level differentially expressed genes (DEGs) related to previously reported fly and rodent models of inter- and/or trans-generational inheritance (**Table S1**). The DEGs represented comparison of stimulated with unstimulated individuals, or of descendants of stimulated with that of unstimulated individuals. With no rat model qualifying the DEG calling criteria used, adjusted p <0.05 and fold change (FC) >1.2, we considered only mouse models for rodent analysis. Altogether, DEGs related to both inter- and trans-generational inheritance in flies, and intergenerational alone in mice formed the compendium. The triggers used in flies included cold exposure (CE), low-protein diet (LPD), restraint stress (RS), and high-sugar diet (HSD) for intergenerational inheritance, and HSD and the centrally acting agent pentylenentetrazole (PTZ) for TEI. The mouse triggers for intergenerational inheritance included CE, LPD, high-fat diet combined with insulin-producing cell (IPC) ablating agent streptozotocin, aging, and phthalate exposure. Besides DEGs, for comparison purpose, we also assembled a set of genome level differentially methylated region (DMR) associated genes in mouse intergenerational models. The DEGs and DMRs were related to diverse tissues (**Table S1**).

The fly DEGs were examined first, with the visual observation suggesting abundance of proteolysis genes (**Fig. 1A**). Indeed, gene ontology (GO) analysis of individual DEG lists showed proteolysis as the most commonly overrepresented biological process term, with a majority of CE and HSD samples displaying enrichment for the term (**Fig. 1B**). This finding was consistent with known regulatory interconnectedness between proteolysis and central metabolic pathways (*6–8*), with cold stress as an energy metabolism modulator. Genome-wide proteolysis genes that were differentially expressed in CE and HSD samples showed only a minor overlap between the models, suggesting that though proteolysis as a biological process is commonly enriched, there exists model specificity at individual gene level (**Fig. 1C**). Next, analysis of TEI data showed that a majority of F0 founder DEGs are oppositely regulated in F2, in both HSD and PTZ models (**Fig. 1D**). To note, this gene expression directionality change was mostly consistent with the phenotypic observation of triglyceride (TAG) level change in F2 in the HSD TEI model, and in a CE TEI model described in a non-transcriptomic study (*5*). Next, clustering based on DEG overlaps among all fly samples showed a trend for positive enrichment in general, with intra-model samples exhibiting higher overlaps than inter-model samples (**Fig. 1E**). This was overall consistent with the commonalities and specificities of models observed above. Further, clustering based on DEG correlation coefficients showed frequent disruption of correlation among samples (**Fig. 1F**). This was also consistent with reversal of gene expression directionality observed above.

**Fig. 1.**
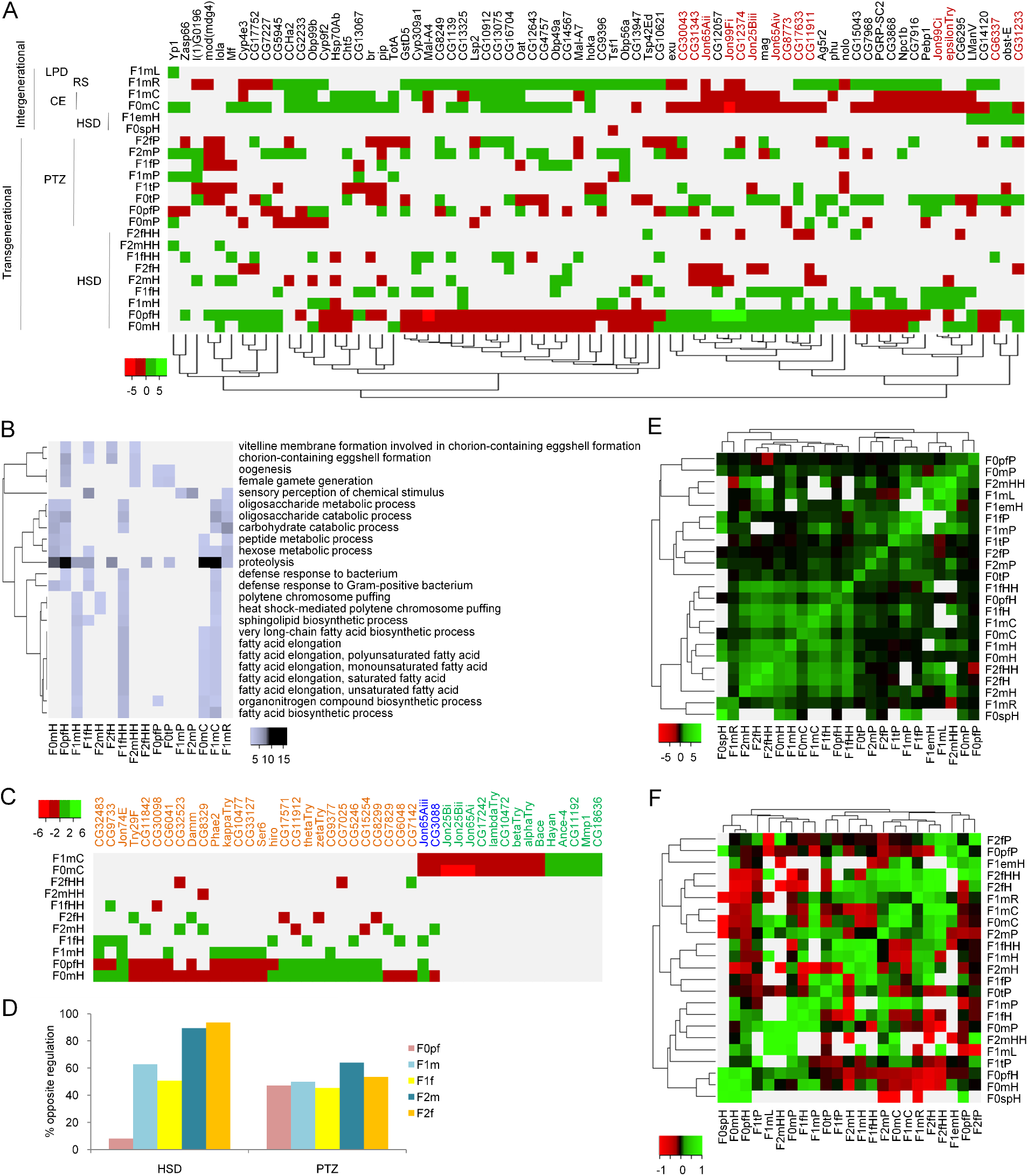
Integrative analysis of previous fly models. (**A**) Heatmap of log_2_FC for most common DEGs among all samples of the intergenerational and TEI models. Number of shades is reduced to discretize gene regulation. Proteolysis genes are indicated in red. Sample labels are explained in **Table S1**. Source tissue is indicated only for sperm (sp), testis (t), and embryo (em). F0pf indicates females not used in cross but treated and handled in parallel with the F0 founder males. (**B**) Heatmap of -log_10_ adjusted p values for most commonly enriched GO terms in complete lists of DEGs. Full set of enriched terms is indicated in **Table S2**. (**C**) Heatmap of log_2_FC for all proteolysis genes in the fly genome except, to avoid duplication, those shown in A. Gene symbols shown in red, green, and blue indicate DEGs unique to HSD and CE, and common to both, in that order. (**D**) Bar chart showing percentage of F0 founder DEGs regulated in opposite direction in rest of the samples. (**E**) Heatmap of log_2_FC in DEG enrichment matrix. Enrichment p values are given in **Table S3**. (**F**) Heatmap based on DEG correlation coefficient matrix.

We next examined mouse DEGs (**Fig. 2A**). Like in fly GO analysis, proteolysis was one of the most commonly enriched terms, with CE, LPD, IPC ablation, and aging models showing the enrichment (**Fig. 2B**). Again, the result was consistent with known association of proteolysis with central metabolic pathways, as mentioned, and aging (*7–9*). Next, like fly, mouse models were mostly associated with specific proteolysis genes (**Fig. 2C**). Regarding directionality of gene expression, comparison of tissue matched samples with sufficient number of DEGs in F0 and F1, available only for CE model, showed largely consistent regulation of genes between the two generations (data not presented), as also visually apparent from the DEG clustering presented above (**Fig. 2A**). To note, maintenance of gene expression directionality between fathers and offspring was found overall consistent with reported phenotypic directionality across various parameters measured in these mouse models. Further, gene set overlap matrix based clustering exhibited, similar to fly analysis, a trend for positive enrichment in general, and a higher enrichment among intra-model than inter-model samples (**Fig. 2D**). In correlation clustering, as expected from the above observations about directionality, correlation was observed among intra-model samples (**Fig. 2E**). Finally, like DEGs, DMRs were also shared among intra-model samples more than inter-model samples (**Fig. 2F**). In contrast, however, GO analysis did not reveal proteolysis enrichment in DMRs (**Fig. 2G**), in any of the sample (**Table S7**).

**Fig. 2.**
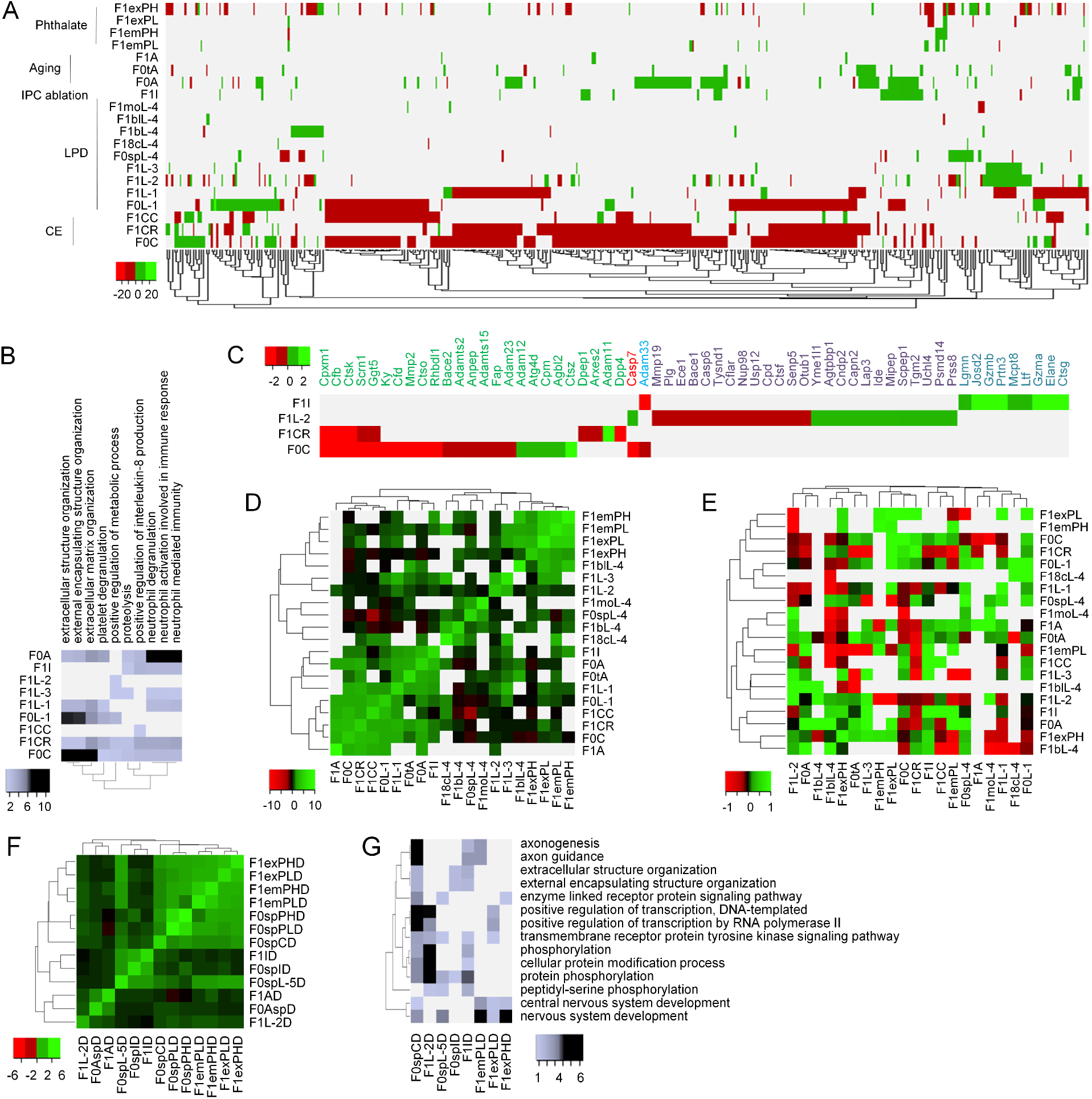
Integrative analysis of previous mouse models. (**A**) Heatmap of 349 most common DEGs. Source tissue is indicated only for sperm (sp), testis (t), embryo (em), 8 cell (8c) embryonic stage, blastula (b), late blastula (bl), morula (mo), and extraembryonic (ex) sample. Numbers in LPD samples indicate independent studies. Sample labels are explained in **Table S1**. (**B**) Heatmap for GO enrichment. Complete set of enriched terms is indicated in **Table S4**. (**C**) Heatmap of all proteolysis genes in the mouse genome except those shown in A. Gene symbols shown in green, purple, aqua, red, and blue indicate DEGs unique to CE, LPD, and IPC ablation, and common to CE and LPD, and CE and IPC ablation models, in that order. (**D**) Heatmap of DEG enrichment matrix. Enrichment p values are given in **Table S5**. (**E**) Heatmap of DEG correlation coefficient matrix. (**F**) Heatmap of DMR enrichment matrix. Enrichment p values are given in **Table S6**. Sample labels are explained in **Table S1**. The last alphabet D in the labels indicates DMRs. (**G**) Heatmap for GO enrichment in DMRs. Complete set of enriched terms is presented in **Table S7**. Other details as correspondingly mentioned in Fig. 1.

The above analysis of previous data suggested association of metabolic trigger induced non-genetic inheritance with proteolysis, in both fly and mouse models. It also suggested phenotypic directionality switching in transgenerational flies. We next turned towards experimentally verifying these observations. As one of the mouse models included IPC ablation induced intergenerational inheritance of prediabetic conditions, and insulin signaling is mostly conserved over animal evolution – with IPC ablation known to induce diabetic-like phenotypes in fly, and mammalian and fly IPCs considered functionally analogous (*10*) – we predicted that this trigger would also induce inter- and trans-generational inheritance in flies. We also predicted that IPC ablation induced inheritance in flies would entail altered expression of genes enriched for proteolysis, with altered proteolysis genes being mostly distinct from other models, and would result in reversal in the regulatory direction of genes affected in the founders in transgenerational individuals. It was also predicted that TAG levels, alterations in which are associated with IPC ablation in flies (*11*), would be affected in the model, with transgenerational flies showing opposite regulation.

The fly IPCs were genetically ablated in F0 males using GAL4/UAS system - wherein the promoter of insulin-like peptide 3 (*ilp3*), that is expressed in IPCs in brain, drives the expression of GAL4 transcriptional activator, and GAL4 inducible UAS enhancer activates the expression of the pro-apoptotic gene reaper (rpr), resulting in constitutive IPC ablation (*12*) – and patrilineal F1-F3 generations obtained, with w^1118^ wild-type (WT) background strain serving as the control lineage (**Fig. 3A**). The precursor strains used for generating the self-regenerating F0 founder line (**Fig. S1**) were outcrossed to w^1118^ for 20 generations, to minimise confounding due to genetic background differences. As presence of transgenic constructs and balancers in F0 and F1 could also potentially confound, the experiment was extended up to F3, not F2, as was the case with above fly TEI models, to ascertain inheritance in WT flies two generations away from non-WT ancestors. Of the four F1 genotypes, the one with both *CyO* and *Ser* balancers and without any of the *ilp3*-GAL4 and UAS-*rpr* constructs was used for obtaining F2 WT, which in turn gave rise to F3 generation (**Fig. 3A**). As IPC ablation effects in *ilp3*-GAL4 x UAS-*rpr* flies have previously been studied using whole body microarray gene expression profiling (*12*), we performed the same to compare all test lineage genotypes, including those three in F1 which were not used in further cross but could have been nonetheless useful in result interpretation, with control lineage flies. The DEGs were identified using the same criteria that were applied above for the analysis of previous models. This resulted in a total of 14 DEG lists, related to *ilp3*-GAL4 x UAS-*rpr* F0 founder males, *ilp3*-GAL4 x UAS-*rpr* females that were not used in cross but were handled in parallel with the F0 founder males, F1 males and females of each of the four genotypes, and F2-F3 WT males and females (**Table S8**). These lists consisted of varying number of DEGs, with 746 being the average number (**Fig. 3B**). As intergenerational comparisons of control lineage males or females yielded an average of only 31 DEGs (data not presented), the above 14 DEG lists were confirmed as largely representing transcriptomic alterations that cannot be ascribed to fly batch or generational effects.

**Fig. 3.**
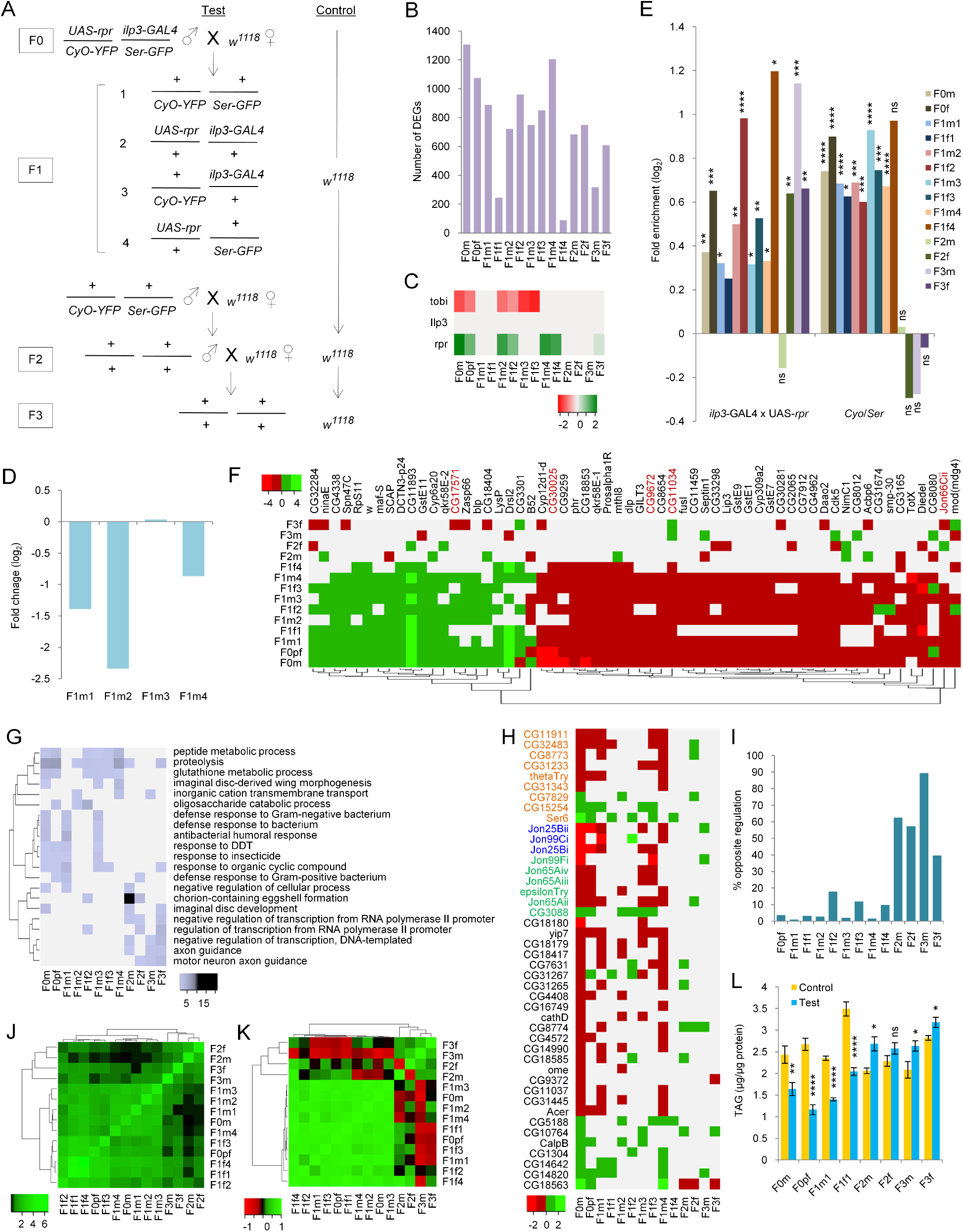
Experimental testing of predictions derived from the analysis of previous models. **(A)** Crossing scheme. (**B**) Bar chart displaying number of DEGs. Sample labels as indicated in A (m, males; f, females), except F0pf, that indicates *ilp3*-GAL4 x UAS-*rpr* females not used in cross but handled in parallel with F0m. Log_2_FC of DEGs and full sample description are presented in **Table S8**. (**C**) Heatmap of log_2_FC showing DEG status. (**D**) Log_2_FC of *ilp2* mRNA in test flies, compared to w^1118^ calibrator. (**E**) Log_2_FC for enrichment of ablation and balancer associated DEGs. (**F**) Most common DEGs. (**G**) Most commonly enriched GO terms. Complete set of enriched terms is indicated in **Table S9**. (**H**) Proteolysis genes in the mouse genome, except those shown in A. Red, blue, and green symbols indicate DEGs common to HSD, CE, and both HSD and CE, in that order. (**I**) Percentage of F0 founder DEGs oppositely regulated in other samples. (**J**) DEG enrichment clustering. Enrichment p values are given in **Table S10**. (**K**) DEG correlation coefficient clustering. (**L**) TAG levels in control and test flies. ns, not significant; hypergeometric (E) and two-tailed t-test (L) p, *<0.05, **<0.01, ***<0.001, and ****<0.0001. Other details as correspondingly mentioned in Fig. 1.

Given that whole fly microarray profiling had previously found in *ilp3*-GAL4 x UAS-*rpr* females extreme downregulation of *tobi*, a functionally validated peripheral target of brain insulin (*12*), we first inspected the status of this gene in our DEGs. As *ilp3* is also expressed peripherally (*13*), we additionally inspected the status of *ilp3* and *rpr* for comparison purpose, assuming ablation, that may not cause absolute loss of cells (*14*), might show up as *ilp3* downregulation and *rpr* upregulation, in case our whole tissue profiling is efficient enough to differentiate transcript abundance between test and control flies. Startlingly, *tobi* was downregulated not only in the ablating genotypes – F0 males, parallel females, and *ilp3*-GAL4 x UAS-*rpr* F1 offspring – but also in non-ablating *ilp3*-GAL4 x Cyo-YFP F1 genotype, and, furthermore, while *ilp3* showed altered regulation in none of the samples, *rpr* was found upregulated in F0-F1 ablating genotypes, non-ablating *UAS-rpr* x Ser-GFP F1 genotype, and F3 WT females (**Fig. 3C**). First, downregulation of *tobi* in non-ablated F1 offspring indicated occurrence of non-genetic inheritance. Next, as GAL4 is required for UAS activation of target gene expression (*15*), *rpr* upregulation in non-ablating F1 genotype and F3 WT flies also suggested intergenerational survival of gene regulatory information. Regarding *ilp3*, altered expression not even in ablating genotypes could have possibly resulted owing to limits in microarray detection. Indeed, given that *ilp2* transcript levels in IPCs exceed that of *ilp3* by almost a degree of magnitude (*10*), *ilp2* was found downregulated in F0 parallel females (**Table S8**). As expected, real-time RT-PCR measurements of *ilp2* RNA abundance in F1 offspring showed decreased levels in three of the four genotypes, with the ablating genotype showing the lowest (**Fig. 3D**). Combined, these results supported inheritance of IPC ablation effects in our model.

Next, we separated inherited and genotypic effects. Gene set overlap test, as described above, was used to examine each of the 14 DEG lists for enrichment of an independent list (*12*) of whole tissue DEGs between IPC ablated, *ilp3*-GAL4 x UAS-*rpr* females and unablated, *ilp3*-GAL4 x UAS-*lacZ* females as well as of another independent list (*16*) of embryonic DEGs comparing allele-specific expression between *CyO/Ser* balancer and wild-type haplotypes (**Table S1**). In order to avoid known gene set size related biases in enrichment analysis, we once deviated from significance and FC criteria set throughout for DEG calling and applied a single criterion of 2.8 FC for the former independent list (*12*) – that pertained to older cDNA microarray technology commonly used for identifying FC based DEGs – so that it becomes equal in size to the latter independent list (*16*), and their respective enrichment in our DEGs can be directly compared. Notably, the female IPC ablation set (*12*) was enriched across F0-F3 in general, with gender matching samples displaying greater overlap, whereas the balancer associated set (*16*) showed enrichment only in F0-F1 (**Fig. 3E**). Together, these results demonstrated occurrence of IPC ablation induced TEI in our model, with F1 showing both inherited and genotypic effects, and F2-F3 only inherited effects.

Finally, we repeated the analysis performed on previous models with our new model (**Fig. 3F**). Gene ontology analysis indeed showed proteolysis as the most common term enriched across F0-F3 samples (**Fig. 3G**). Also, as expected, the new model was associated with proteolysis genes mostly distinct from those related to previous TEI models (**Fig. 3H**). Further, as predicted, the F0 founder genes were largely regulated in the same direction in F1, and in the opposite direction in F2-F3 (**Figure 3I**). Next, gene set overlap clustering separated F0-F1 from F2-F3, consistent with genotypic differences between inter- and trans-generational flies (**Fig. 3J**). Also, consistent with gene regulatory differences observed above, DEG correlation clustering separated F0-F1 from F2-F3, with the samples in the latter group showing a trend for loss of correlation with those in the former (**Fig. 3K**). Lastly, as predicted, whole body TAG levels under HSD condition, a drastic decrease in which is known in *ilp3*-GAL4 x UAS-*lacZ* flies (*11*), showed a decrease in F0 males, parallel females, and F1 offspring, and a trend for increase in F2-F3 (**Fig. 3L**). Combined, predictions based on analysis of previous data were experimentally confirmed in flies.

## Conclusion

We have shown here that transcriptomic responses in non-genetic inheritance are interchangeable between fly and mouse. This uplifts the plausibility of mammalian TEI to elevated levels. We have also shown that phenotypic directionality changes as the inheritance progresses from inter- to trans-generational stage. This suggests that non-genetic inheritance may have both adaptive and maladaptive consequences, depending on presence or absence of ancestral environmental conditions. The observed cross-species functional conservation and cross-generational phenotypic directionality shift together supports evolutionary and physiological implications of epigenetic inheritance. Our consolidated hypothesis forming and hypothesis testing results provide a sound basis for spurring future evolvability and mechanistic studies in ancestral memory propagation across generations.

## Materials and Methods

### DEGs and DMRs from previous studies

Available studies were searched in NCBI’s PubMed and Gene Expression Omnibus (GEO) using relevant key words, related to paternal exposure and patrilineal models of intergenerational inheritance and TEI in *Drosophila*, mice, and rats, as also IPC ablation and balancers in flies. The DEGs were identified from publications, when provided in full, or from GEO accessions, directly or through GEO2R, CARMAweb, and GREIN analysis for oligonucleotide microarray, cDNA microarray, and RNA-seq, in that order (*17–19*). The DEGs were uniformly identified using the criteria of, unless stated otherwise, 0.05 adjusted p value and 1.2 FC cut-offs. No mouse model of transgenerational, and no rat model of either inter- or trans-generational inheritance qualified these criteria for inclusion. Therefore, fly models of both inter- and trans-generational, and mouse models of only intergenerational inheritance were finally considered in the analysis. In non-genetic inheritance, the DEGs identified represented comparison between stimulated and unstimulated individuals, or between descendants of stimulated and unstimulated individuals. Similarly, the DMR genes as reported in on-genetic inheritance were directly used. The DEGs and DMR genes were normalized against 12,858 FlyMine or 23,353 MouseMine gene symbols (*20, 21*) representing whole fly or whole mouse genome, in that order. FC values of DEGs were expressed on a logarithmic scale to base 2.

The DEG compendium for non-genetic inheritance comprised of 23 lists belonging to a total of 6 individually reported fly models (*4*, *22–26*), and 20 lists belonging to a total of 8 individually reported mouse models (*27–34*). The source tissues of the fly DEGs included sperm, testis, embryo, whole fly, and fly head or other body parts. The source tissues of the mouse DEGs included sperm, testis, embryo, extraembryonic tissue, liver, brown adipose tissue, white adipose tissue, pancreatic islets, and hippocampus. The DMR genes were related to several of the above mouse models as well as an additional LPD model that belonged to a non-transcriptomic study (*27–31, 35*). Overall, the DMR gene lists, 13 in number, were associated with a total of 6 individually reported models. The source tissue of the DMRs included sperm, embryo, extraembryonic tissue, liver, pancreatic islets, and brain.

DEGs related to previous IPC ablation study in flies represented comparison of ablated female flies with corresponding non-ablated controls (*12*). DEGs associated with fly balancers represented allele-specific embryonic expression between balancer and wild-type haplotypes (*16*).

The log_2_FC values associated with the DEGs, the lists of DMR genes, and genomic background genes used in the present analysis, along with references and details of the reported studies, and labels representing them in the figures in the present work, are presented in **Table S1**.

### Heatmaps and clustering

Heatmaps were generated using Heatmapper (*36*), with, as applicable, the scale type none, the clustering method average linkage, and the distance measurement method Pearson. Other settings included Red/Green or custom color scheme, 50 color brightness, and for all except DEG heatmaps, 100 number of shades. For DEG heatmaps, the number of shades was reduced to 4, to discretize up- and down-regulation for visualization.

### Gene ontology

Gene ontology analysis was performed using FlyEnrichr or Enrichr, as applicable, to identify biological process terms enriched in DEG and DMR gene lists (*37*). Adjusted p value cut-off of 0.05 was used for identifying enriched terms. Proteolysis genes in fly and mouse genomes were identified from the gene ontology resource (*38*).

### Gene regulation directionality

For calculating directionality of gene regulation, genes differentially expressed in F0 founder males were examined for their DEG status in identical tissues in other F0-F3 samples, as applicable. Genes showing regulation opposite of that in the founders in a given non-founder sample were counted and expressed in terms of percentage of total number of DEGs in that sample.

### Gene set overlap

Gene set overlap was calculated as fold enrichment between two samples in the whole genome background of aforementioned 12,858 fly or 23,353 mouse genes, as applicable. A p value cutoff of 0.05 in hypergeometric test, with sampling without replacement from a finite population, was used to call significant overlaps.

### Gene expression correlation

Correlation coefficients of differential gene expression were obtained using log_2_FC values for all the DEGs in a given sample set.

### Fly husbandry

Standard methods of fly handling and manipulation were used. Flies were maintained under artificial light-dark cycles of 12 hr each, with lights on at 9:00 hr. Rearing temperature was 25±1^0^C. A previously described fly food was used, with the food consisting of yeast extract, sugar, peptone, dry yeast powder, agar, propionic acid, and methyl 4-hydroxybenzoate (*4*).

### Fly genotypes

The *ilp3*-GAL4 line was a gift from Michael Pankratz. The w^1118^ (#3605), *UAS-rpr* (#5824), CyO-YFP (#8578), and Ser-GFP (#4534) lines were obtained from the Bloomington Drosophila Stock Center. The *CyO, TM6B*, and *Sb* lines are available widely and were used as is. Routine fly genetic methods were employed. The stable inheritance of transgenic constructs was confirmed in the F1 generation resulting from the backcross of the F0 founders with w^1118^, with the phenotypes occurring in predicted Mendelian ratio. Crosses were set up following confirmation of appropriate phenotypic and fluorescence markers.

### Real time RT-PCR

The template RNA was extracted by homogenizing 80-90 male flies suspended in TRIzol™ (ThermoFisher, #15596026) in a Mini-Beadbeater-16 (BioSpec, #607EUR) containing 0.7 mm Zirconia beads (#11079107zx). Real-time RT-PCR was performed to assess the levels of *ilp2* (TaqMan assay ID Dm01822534_g1) in the test lineage F1 genotypes, with *RpL32* (Dm02151827_g1) serving as the reference gene and the control lineage w^1118^ as the calibrator sample. TB Green® Premix Ex Taq™ (Takara Bio, # RR420L) and LightCycler® 480 Multiwell Plate 384 (Roche, # 04729749001) were used according to the manufacturers’ recommendations. The ΔΔCt method was used to calculate the RNA FC in the test samples, compared to the baseline of the calibrator, using 4-7 biological replicates.

### Gene expression profiling

Flies were homogenized in TRIzol, and the RNA isolated using RNeasy Microarray Tissue Mini Kit (Qiagen, #73304). Amplification and biotin labeling of RNA was performed using GeneChip 3’ IVT express kit (Affymetrix, #901229). Affymetrix Drosophila Genome 2.0 array (# 900533) was used for hybridization. The hybridized arrays were processed using Affymetrix GeneChip Hybridization Wash and Stain Kit (# 900720). The profiling was performed using three to four biological replicates. The data sets were analyzed using the Bioconductor suite for R and the limma package (version 3.46.0). Probe intensities were corrected for background hybridization using GC-RMA, and quantile normalized individually within following sets of test and control arrays: F0 test and F1 control males (3 replicates each); F0 test and F1 control females (3 replicates each); F1 test and F0 control males, with all four test line genotypes separately compared with the common controls (3 replicates each); F1 test and F0 control females, with all four test line genotypes separately compared with the common controls (3 replicates each); F2 test and F2 control males (4 replicates each); F2 test and F2 control females (3 replicates each); F3 test and F3 control males (4 replicates each); F3 test and F3 control females (4 replicates each). DEGs were identified using the aforesaid criteria of FC >1.2 and adjusted p <0.05. The above mentioned 12,858 FlyMine gene symbols (*20*) served as genomic background.

### Triglyceride and protein

Batches of 12 male or 10 female files, suspended in sterile water, were homogenized in a Mini-Beadbeater-16 containing 0.7 mm Zirconia beads, as above, and as described previously (*4*). The homogenates were heat denatured, filtered, and processed for triglyceride (TAG) and protein estimation, using 4-30 biological replicates. LiquiColor™ Triglycerides Kit (Stanbio, #SB2200225) was used for measuring TAG concentrations. Pierce™ BCA Protein Assay Kit (Thermo Fisher Scientific, #23227) was used for determining protein concentrations. Manufacturer recommended protocols were followed.

## Acknowledgements

The work was supported by funding to AS from the BSC0122 network project of the Council of Scientific and Industrial Research, India, and a senior research fellowship to AB by the Indian Council of Medical Research.

## Authors contributions

AB performed the experiments, and analyzed the experimental data. AS conceptualized the study, performed integrative data analysis and bioinformatics, and wrote the manuscript. Both the authors approved the final manuscript.

## Data availability

The microarray raw data will be publicly available in due course.

## Supplementary Information

**Fig. S1**

**Crossing scheme for generating the founder line**. The precursor lines in F1 and F2 were outcrossed with w^1118^ for 20 generations. The F5 represents the founder line, the males of which were used as F0 for investigating transgenerational inheritance (**Fig. 3A**). The stability of the line was checked by performing Mendelian test crosses.

---

**Table S1**

Genomic background genes and gene lists related to previous studies

**Table S2**

Gene ontology enrichment in DEGs associated with previously described fly models

**Table S3**

DEG overlap matrix associated p values for previously described fly models

**Table S4**

Gene ontology enrichment in DEGs associated with previously described mouse models

**Table S5**

DEG overlap matrix associated p values for previously described mouse models

**Table S6**

DMR overlap matrix associated p values for previously described mouse models

**Table S7**

Gene ontology enrichment in DMR genes associated with previously described mouse models

**Table S8**

DEGs associated with the newly developed IPC ablation fly model

**Table S9**

Gene ontology enrichment in DEGs associated with the newly developed IPC ablation fly model

**Table S10**

DEG overlap matrix associated p values for the newly developed IPC ablation fly model

## References

1. G. Cavalli, E. Heard, Advances in epigenetics link genetics to the environment and disease. Nature 571, 489–499 (2019).

2. M. H. Fitz-James, G. Cavalli, Molecular mechanisms of transgenerational epigenetic inheritance. Nat. Rev. Genet. 23, 325–341 (2022).

3. M. F. Perez, B. Lehner, Intergenerational and transgenerational epigenetic inheritance in animals. Nat. Cell Biol. 21, 143–151 (2019).

4. M. Teltumbade, A. Bhalla, A. Sharma, Paternal inheritance of diet induced metabolic traits correlates with germline regulation of diet induced coding gene expression. Genomics 112, 567–573 (2020).

5. P. Karunakar, A. Bhalla, A. Sharma, Transgenerational inheritance of cold temperature response in Drosophila. FEBS Lett. 593, 594–600 (2019).

6. R. T. Brookheart, J. G. Duncan, Modeling dietary influences on offspring metabolic programming in Drosophila melanogaster. Reproduction 152, R79–90 (2016).

7. F. Ottens, A. Franz, T. Hoppe, Build-UPS and break-downs: metabolism impacts on proteostasis and aging. Cell Death Differ. 28, 505–521 (2021).

8. L. H. Manfredi, N. M. Zanon, M. A. Garófalo, L. C. Navegantes, I. C. Kettelhut, Effect of short-term cold exposure on skeletal muscle protein breakdown in rats. J. Appl. Physiol. (1985) 115, 1496–1505 (2013).

9. D. N. Reeds, W. T. Cade, B. W. Patterson, W. G. Powderly, S. Klein, K. E. Yarasheski, Whole-body proteolysis rate is elevated in HIV-associated insulin resistance. Diabetes 55, 2849–2855 (2006).

10. J. Cao, J. Ni, W. Ma, V. Shiu, L. A. Milla, S. Park, M. L. Spletter, S. Tang, J. Zhang, X. Wei, S. K. Kim, M. P. Scott, Insight into insulin secretion from transcriptome and genetic analysis of insulin-producing cells of Drosophila. Genetics 197, 175–192 (2014).

11. S. N. Al Saud, A. C. Summerfield, N. Alic, Ablation of insulin-producing cells prevents obesity but not premature mortality caused by a high-sugar diet in Drosophila. Proc. Biol. Sci. 282, 20141720 (2015).

12. S. Buch, C. Melcher, M. Bauer, J. Katzenberger, M. J. Pankratz, Opposing effects of dietary protein and sugar regulate a transcriptional target of Drosophila insulin-like peptide signaling. Cell Metab. 7, 321–332 (2008).

13. L. E. O’Brien, S. S. Soliman, X. Li, D. Bilder, Altered modes of stem cell division drive adaptive intestinal growth. Cell 147, 603–614 (2011).

14. S. J. Broughton, M. D. Piper, T. Ikeya, T. M. Bass, J. Jacobson, Y. Driege, P. Martinez, E. Hafen, D. J. Withers, S. J. Leevers, L. Partridge, Longer lifespan, altered metabolism, and stress resistance in Drosophila from ablation of cells making insulin-like ligands. Proc. Natl. Acad. Sci. U.S.A. 102, 3105–3110 (2005).

15. C. B. Phelps, A. H. Brand, Ectopic gene expression in Drosophila using GAL4 system. Methods 14, 367–379 (1998).

16. Y. Ghavi-Helm, A. Jankowski, S. Meiers, R. R. Viales, J. O. Korbel, E. E. M. Furlong, Highly rearranged chromosomes reveal uncoupling between genome topology and gene expression. Nat. Genet. 51, 1272–1282 (2019).

17. T. Barrett, S. E. Wilhite, P. Ledoux, C. Evangelista, I. F. Kim, M. Tomashevsky, K. A. Marshall, K. H. Phillippy, P. M. Sherman, M. Holko, A. Yefanov, H. Lee, N. Zhang, C. L. Robertson, N. Serova, S. Davis, A. Soboleva A, NCBI GEO: archive for functional genomics data sets--update. Nucleic Acids Res. 41(Database issue), D991–D995 (2013).

18. J. Rainer, F. Sanchez-Cabo, G. Stocker, A. Sturn, Z. Trajanoski, CARMAweb: comprehensive R-and Bioconductor-based web service for microarray data analysis Nucleic Acids Res. 34(Web Server Issue), W498–503 (2006).

19. N. A. Mahi, M. F. Najafabadi, M. Pilarczyk, M. Kouril, M. Medvedovic, GREIN: An Interactive Web Platform for Re-analyzing GEO RNA-seq Data. Sci Rep. 9, 7580 (2019).

20. R. Lyne, R. Smith, K. Rutherford, M. Wakeling, A. Varley, F. Guillier, H. Janssens, W. Ji, P. Mclaren, P. North, D. Rana, T. Riley, J. Sullivan, X. Watkins, M. Woodbridge, K. Lilley, S. Russell, M. Ashburner, K. Mizuguchi, G. Micklem, FlyMine: an integrated database for Drosophila and Anopheles genomics. Genome Biol. 8, R129 (2007).

21. H. Motenko, S. B. Neuhauser, M. O’Keefe, J. E. Richardson, MouseMine: a new data warehouse for MGI. Mamm. Genome 26, 325–330 (2015).

22. A, Öst, A. Lempradl, E. Casas, M. Weigert, T. Tiko, M. Deniz, L. Pantano, U. Boenisch, P. M. Itskov, M. Stoeckius, M. Ruf, N. Rajewsky, G. Reuter, N. Iovino, C. Ribeiro, M. Alenius, S. Heyne, T. Vavouri, J. A. Pospisilik, Paternal diet defines offspring chromatin state and intergenerational obesity. Cell 159, 1352–1364 (2014).

23. F. Zajitschek, S. Zajitschek, M. Manier, Paternal diet affects differential gene expression, but not sperm competition, in sons. Biol. Lett. 13, 20160914 (2017).

24. A. Zare, A. M. Johansson, E. Karlsson, N. Delhomme, P. Stenberg, The gut microbiome participates in transgenerational inheritance of low-temperature responses in Drosophila melanogaster. FEBS Lett. 592, 4078–4086 (2018).

25. K. H. Seong, N. H. Ly, Y. Katou, N. Yokota, R. Nakato, S. Murakami, A. Hirayama, S. Fukuda, S. Kang, T. Soga, K. Shirahige, S. Ishii, Paternal restraint stress affects offspring metabolism via ATF-2 dependent mechanisms in Drosophila melanogaster germ cells. Commun. Biol. 4, 208 (2020).

26. A. Sharma, P. Singh, Detection of transgenerational spermatogenic inheritance of adult male acquired CNS gene expression characteristics using a Drosophila systems model. PLoS One 4, e5763 (2009).

27. W. Sun, H. Dong, A. S. Becker, D. H. Dapito, S. Modica, G. Grandl, L. Opitz, V. Efthymiou, L. G. Straub, G. Sarker, M. Balaz, L. Balazova, A. Perdikari, E. Kiehlmann, S. Bacanovic, C. Zellweger, D. Peleg-Raibstein, P. Pelczar, W. Reik, I. A. Burger, F. von Meyenn, C. Wolfrum, Cold-induced epigenetic programming of the sperm enhances brown adipose tissue activity in the offspring. Nat. Med. 24, 1372–1383 (2018).

28. Y. Wei, C. R. Yang, Y. P. Wei, Z. A. Zhao, Y. Hou, H. Schatten, Q. Y. Sun, Paternally induced transgenerational inheritance of susceptibility to diabetes in mammals. Proc. Natl. Acad. Sci. U.S.A. 111, 1873–1878 (2014).

29. B. R. Carone, L. Fauquier, N. Habib, J. M. Shea, C. E. Hart, R. Li, C. Bock, C. Li, H. Gu, P. D. Zamore, A. Meissner, Z. Weng, H. A. Hofmann, N. Friedman, O. J. Rando, Paternally induced transgenerational environmental reprogramming of metabolic gene expression in mammals. Cell 143, 1084–1096 (2010).

30. K. Xie, D. P. Ryan, B. L. Pearson, K. S. Henzel, F. Neff, R. O. Vidal, M. Hennion, I. Lehmann, M. Schleif, S. Schröder, T. Adler, B. Rathkolb, J. Rozman, A. L. Schütz, C. Prehn, M. E. Mickael, M. Weiergräber, J. Adamski, D. H. Busch, G. Ehninger, A. Matynia, W. S. Jackson, E. Wolf, H. Fuchs, V. Gailus-Durner, S. Bonn, M. Hrabé de Angelis, D. Ehninger, Epigenetic alterations in longevity regulators, reduced life span, and exacerbated aging-related pathology in old father offspring mice. Proc. Natl. Acad. Sci. U.S.A. 115, E2348–E2357 (2018).

31. O. A. Oluwayiose, C. Marcho, H. Wu, E. Houle, S. A. Krawetz, A. Suvorov, J. Mager, J. Richard Pilsner. Paternal preconception phthalate exposure alters sperm methylome and embryonic programming. Environ. Int. 155, 106693 (2021).

32. N. H. Ly, T. Maekawa, K. Yoshida, Y. Liu, M. Muratani, S. Ishii, RNA-Sequencing Analysis of Paternal Low-Protein Diet-Induced Gene Expression Change in Mouse Offspring Adipocytes. G3 (Bethesda) 9, 2161–2170 (2019).

33. K. Yoshida, T. Maekawa, N. H. Ly, S. I. Fujita, M. Muratani, M. Ando, Y. Katou, H. Araki, F. Miura, K. Shirahige, M. Okada, T. Ito, B. Chatton, S. Ishii, ATF7-Dependent Epigenetic Changes Are Required for the Intergenerational Effect of a Paternal Low-Protein Diet. Mol. Cell 78, 445–458.e6 (2020).

34. U. Sharma, C. C. Conine, J. M. Shea, A. Boskovic, A. G. Derr, X. Y. Bing, C. Belleannee, A. Kucukural, R. W. Serra, F. Sun, L. Song, B. R. Carone, E. P. Ricci, X. Z. Li, L. Fauquier, M. J. Moore, R. Sullivan, C. C. Mello, M. Garber, O. J. Rando, Biogenesis and function of tRNA fragments during sperm maturation and fertilization in mammals. Science 351, 391–396 (2016).

35. Watkins AJ, Dias I, Tsuro H, Allen D, Emes RD, Moreton J, Wilson R, Ingram RJM, Sinclair KD. Paternal diet programs offspring health through sperm-and seminal plasma-specific pathways in mice. Proc. Natl. Acad. Sci. U.S.A. 115, 10064–10069 (2018).

36. S. Babicki, D. Arndt, A. Marcu, Y. Liang, J. R. Grant, M. Maciejewski, D. S. Wishart, Heatmapper: web-enabled heat mapping for all. Nucleic Acids Res. 44(W1), W147–W153 (2016).

37. Z. Xie, A. Bailey, M. V. Kuleshov, D. J. B. Clarke, J. E. Evangelista, S. L. Jenkins, A. Lachmann, M. L. Wojciechowicz, E. Kropiwnicki, K. M. Jagodnik, M. Jeon, A. Ma’ayan, Gene Set Knowledge Discovery with Enrichr. Curr. Protoc. 1, e90 (2021).

38. Gene Ontology Consortium. The Gene Ontology resource: enriching a GOld mine. Nucleic Acids Res. 49(D1), D325–D334 (2021).

