## Supplementary Figure 1 for "Phenotypic cross-species conservation and cross-generation directionality switching in epigenetic inheritance"

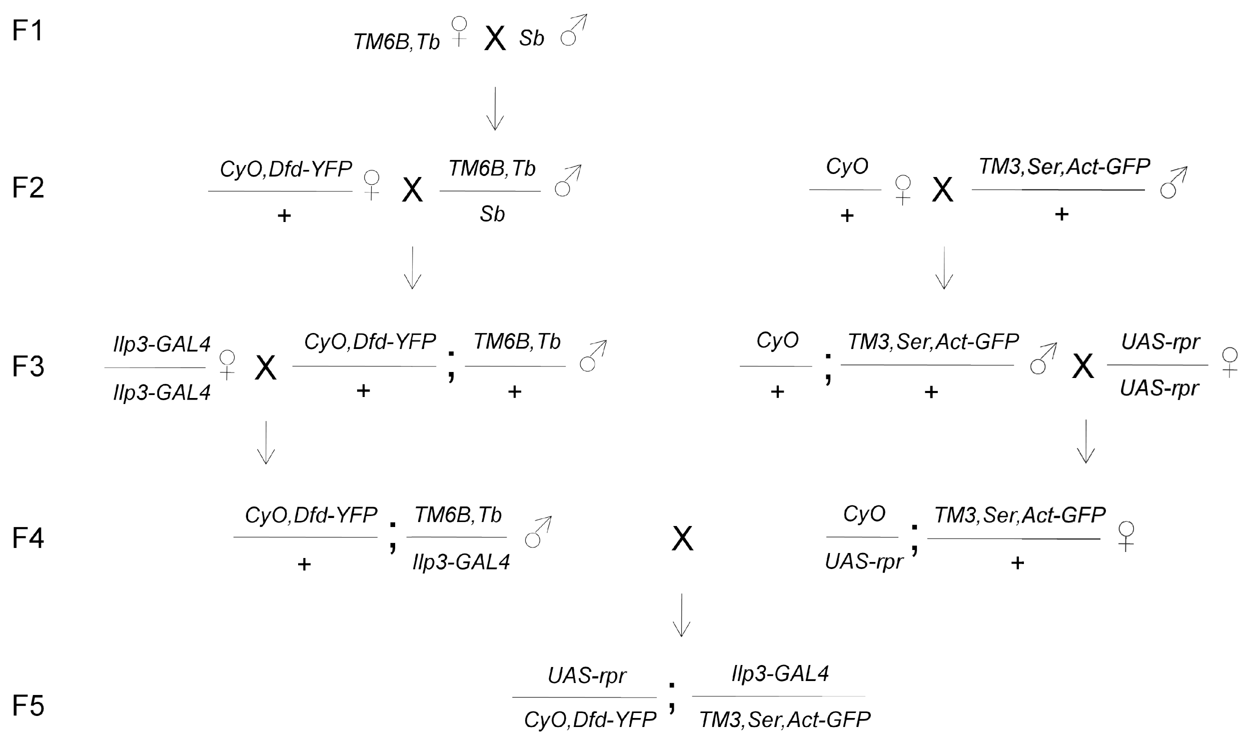


**Fig. S1. Crossing scheme for generating the founder line**. The precursor lines in F1 and F2 were outcrossed with $w^{1118}$ for 20 generations. The F5 represents the founder line, the males of which were used as F0 for investigating transgenerational inheritance (**Fig. 3A**). The stability of the line was checked by performing Mendelian test crosses.
